# A high-density diffuse optical tomography dataset of naturalistic viewing

**DOI:** 10.1101/2023.11.07.565473

**Authors:** Arefeh Sherafati, Aahana Bajracharya, Michael S. Jones, Emma Speh, Monalisa Munsi, Chen-Hao P. Lin, Andrew K. Fishell, Tamara Hershey, Adam T. Eggebrecht, Joseph P. Culver, Jonathan E. Peelle

## Abstract

Traditional laboratory tasks offer tight experimental control but lack the richness of our everyday human experience. As a result, many cognitive neuroscientists have been motivated to adopt experimental paradigms that are more natural, such as stories and movies. Here we describe data collected from 58 healthy adult participants (aged 18–76 years) who viewed 10 minutes of a movie (*The Good, the Bad, and the Ugly*, 1966). Most (36) participants viewed the clip more than once, resulting in 106 sessions of data. Cortical responses were mapped using high-density diffuse optical tomography (first-through fourth nearest neighbor separations of 1.3, 3.0, 3.9, and 4.7 cm), covering large portions of superficial occipital, temporal, parietal, and frontal lobes. Consistency of measured activity across subjects was quantified using intersubject correlation analysis. Data are provided in both channel format (SNIRF) and projected to standard space (NIfTI) using an atlas-based light model. These data are suitable for methods exploration as well as investigating a wide variety of cognitive phenomena.

## Background & Summary

Most cognitive neuroscientists are interested in how human brains interact with the real world. To do so, we frequently create tightly controlled laboratory paradigms intended to isolate one or more aspects of sensory or cognitive processing. We hope that the findings from these purposefully artificial experiments will generalize to real-world scenarios. However, is such an assumption justified? To answer this question, <u>and bolstered by methodological advances,</u>cognitive neuroscience has been increasingly moving towards the use of naturalistic stimuli^1–4^.

For over 20 years, researchers using fMRI have explored the use of movies to provide rich stimulation for research participants including children^5–7^, healthy adults^8–13^, and numerous other populations^14–17^. Although not completely naturalistic, movies convey complex auditory and visual information covering a range of sensory, cognitive, and linguistic domains. Movies therefore provide researchers the opportunity to study processing that more closely mimics everyday experience than standard laboratory tasks. Additionally, the ability to concurrently address multiple domains of sensory and cognitive processing may make movies a more efficient way to collect data than traditional cognitive psychology paradigms. For example, 10 minutes of movie data collection might replace 30 minutes of domain-specific data collection (e.g., 10 minutes of an auditory task, 10 minutes of a visual task, 10 minutes of a language task). <u>Movies also facilitate functional connectivity analysis which can identify brain networks based on temporal correspondence.</u>

Optical neuroimaging, particularly functional near infrared spectroscopy (fNIRS), offers many advantages for studying human brain function. First, it is acoustically silent, avoiding the auditory confounds present in fMRI^18^. Second, implanted medical devices are not contraindicated, meaning optical imaging can be used on people with implanted medical devices, such as cochlear implants^19,20^ or implanted electrodes used for deep brain stimulation^21^. Finally, fNIRS facilitates real-world applications including imaging during face-to-face interaction^22^. These advantages make fNIRS well suited for studying cognition in context^23^ that includes social interaction^24^.

Traditionally, drawbacks associated with fNIRS include limited coverage of the cortex, uneven sensitivity over the field of view, and lack of depth information necessary for removing superficial (i.e., non-brain) hemodynamic components. These challenges motivated the development of high-density diffuse optical tomography (HD-DOT), in which a lattice of closely-spaced sources and detectors provides homogenous sensitivity and spatial resolution comparable to that obtained in fMRI (**Figure 1**)^21^.

**Figure 1.**
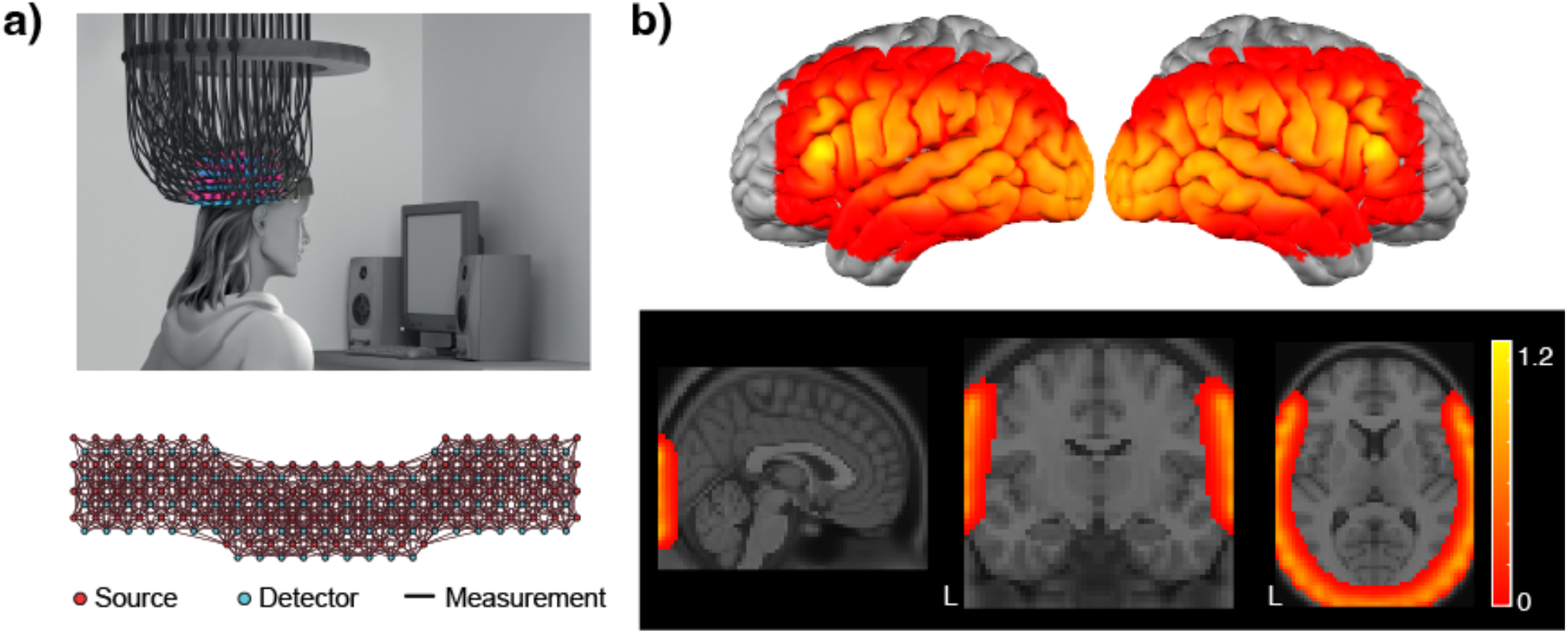
**a)** Illustration of HD-DOT cap and optode arrangement (see ref. 21 for details). **b)** Illustration of field of view in the reconstructed images.

Here, we present data collected using custom HD-DOT during movie-viewing, which complements existing open fMRI data sets of movie viewing^13,25–27^. Although the use of movies in HD-DOT has been previously explored^28^, our hope is that by making the current data available we will facilitate methodological and theoretical advances related to naturalistic stimulation. These include identification of quality control metrics, how to best handle estimated motion, and optimal methods for modeling quasi-continuous variables.

## Method

### Participants

Participants were recruited from the Washington University in Saint Louis community. Data from 58 participants are included in the data set, ranging in age from 18–76 years (M = 23; F = 35; mean age = 31 years); self-reported handedness (right = 55, left = 2, ambidextrous = 1). Forty-nine of the participants reported English as the first language they learned. In addition, participants reported to have normal vision and hearing and no known history of neurological disorders. All participants gave written informed consent prior to the experiment session, which was approved by and carried out in accordance with the Human Research Protection Office at Washington University (protocol numbers 201101896, 201908154, and 202108047).

### Stimulus

The stimulus was a clip of approximately 10 minutes taken from *The Good, the Bad, and the Ugly* (1966), a movie previously used in fMRI^8^ and HD-DOT^28^. All participants viewed between 16:48 and 27:30. A subset of the participants viewed a longer clip; in this case, their data were truncated so that all participants have the same amount of data covering an identical portion of the movie. Copyright restrictions preclude openly sharing the stimulus but guidance is available on request from the corresponding author. We did not systematically document whether participants had previously viewed the selected clip; anecdotally, most participants reported being unfamiliar with the movie.

The same movie clip was used in several projects between 2012–2022 and thus was presented along with various other tasks, including auditory, visual, motor, language, and collection of resting state data. Analyses including 7 of these participants have been published previously^28^ but none of the data is included in a public data set.

### Procedure

Participants were seated on a comfortable chair in an acoustically isolated room facing an LCD screen located 76 cm from them, at approximately eye level, which displayed the movie. The soundtrack was presented through two speakers located approximately 150 cm away at about ± 21° from the subjects’ ears. The sound level was approximately 65 dBA but not calibrated. The HD-DOT cap was fitted to the subject’s head to maximize optode-scalp coupling, assessed via real-time coupling coefficient readouts using in-house software. The movie was presented using Psychophysics Toolbox 3^29^ (RRID SCR_002881) in MATLAB.

### Data acquisition

Data was acquired using two different continuous wave HD-DOT caps. The first cap had 96 sources and 92 detectors (LED; 750 nm and 850 nm)^21^ and was used in 9 participants. <u>This device has been previously described in detail, and been validated against fMRI for mapping localizer tasks, movie viewing, and resting state functional connectivity</u>^21^. The second cap was built by attaching an additional pad on top of the previous cap and had 128 sources and 125 detectors (Laser diode; 685 nm and 830 nm) and was used in the remaining 49 participants. Source-detector pairs were arranged on both caps to enable first-through fourth nearest neighbor separations of 1.3, 3.0, 3.9, and 4.7 cm, respectively. Our in-house software controlled temporal, frequency, and spatial encoding patterns which achieved an overall framerate of 10 Hz.

Full measurement sets are available in the raw data formats (NeuroDOT-compatible.mat files and SNIRF). In order to make the analyses in this paper consistent, we have removed all the measurements from the additional motor pad in 49 participants that were scanned using the bigger cap to contain the same 96 sources and 92 detectors as the other subjects in the preprocessed data formats (NIfTI and BIDS) and all the analyses presented in this paper.

### Data processing

Data processing is schematically illustrated in **Figure 2**. Data preprocessing was done using the NeuroDOT toolbox (https://www.nitrc.org/projects/neurodot) based on the principles of modeling light emission, diffusion, and detection through the head^20,30^. Data processing steps included taking the log-mean ratio of the light levels for each time-point and the temporal mean across the run (as a baseline value). This step was followed by excluding any source-detector measurement that had a temporal standard deviation of 7.5% in the least-noisy 60 sec (lowest mean GVTD)^31^ of each run or higher to exclude any noisy measurements due to poor optode-scalp coupling or movement. In summary, the percentage of measurements remained for each source-detector separation in this dataset was (mean ± STD): 99 ± 1% out of 644 first nearest-neighbor pairs, 95 ± 5% out of 1068 second nearest-neighbor pairs. Due to low SNR, we did not use the measurements beyond second nearest neighbors. The measurements were then high-pass filtered (0.02 Hz cutoff) to remove low-frequency drift. We then estimated and regressed the global superficial signal as the average of all first nearest neighbor measurements (1.3 cm source-detector pair separation)^32^. Following that, data were low-pass filtered to 0.5 Hz cutoff to remove cardiac oscillations. These frequencies were chosen in all previous studies that were published using the existing HD-DOT devices and are consistent with best practices^33^. Data were downsampled from 10 Hz to 1 Hz and then used for image reconstruction.

**Figure 2.**
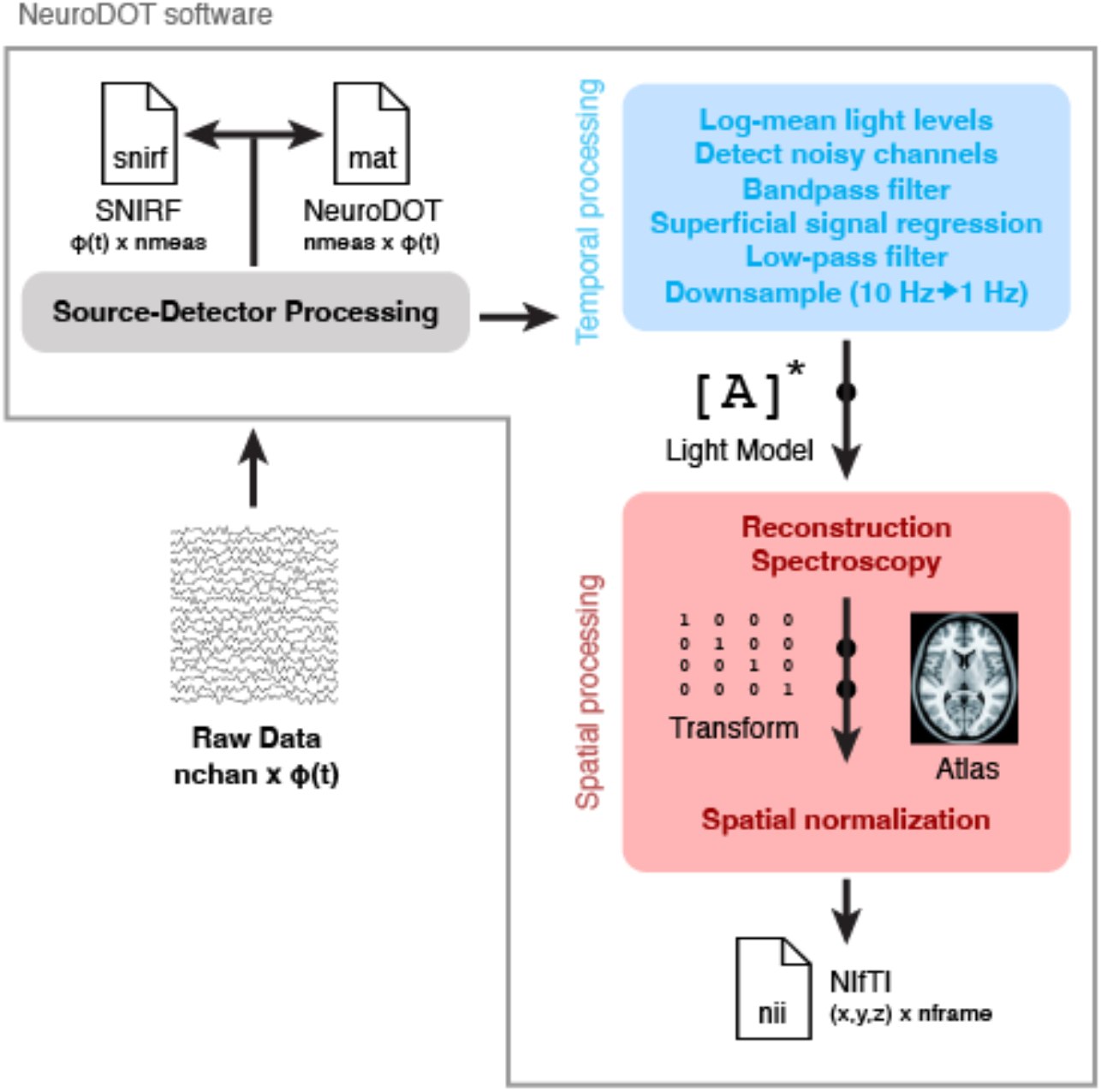
Schematic of data processing and file formats. The data are provided in three different formats: SNIRF, NeuroDOT, and NIfTI.

We then computed a forward model of light propagation based on the two wavelengths used for each device on an anatomical atlas including the non-uniform tissue structures: scalp, skull, CSF, gray matter, and white matter^34^. The resulting sensitivity matrix was then inverted for calculating the relative changes in absorption at the two wavelengths via reconstruction using Tikhonov regularization and spatially variant regularization^21^. Relative changes in oxygenated, deoxygenated, and total hemoglobin (ΔHbO, HbR, ΔHbT) were then computed using the absorption and extinction coefficients of oxygenated and deoxygenated hemoglobin at the two wavelengths. We resampled all data to a 3 × 3 × 3 mm standard atlas using a linear affine transformation.

The preprocessed data were then converted to the NIfTI file format for analysis and sharing purposes. Intersubject correlation analysis was performed using the automatic analysis (aa) environment^35^, version 5.8. Additionally, the *ndot2snirf* function in NeuroDOT was used to convert the raw data to SNIRF^36^ file format followed by *snirf2bids* function to generate other necessary metadata files to satisfy the BIDS specification for NIRS^36,37^. In addition to using these standard functions, the NIfTI files were cropped to align with the actual start and end points of the movie stimulus.

### Data Records

The data records are organized following the Brain Imaging Data Structure (BIDS standard^37^ and are available as dataset ds004569^38^ on OpenNeuro^39^.

### Technical Validation

#### Data quality

To quantify data quality across sessions we focused on two measures intended to capture effects of participant motion and light levels (**Figure 3**). Because we do not have objective measures of motion (e.g., photometry or accelerometers), we rely on a signal-based proxy for motion: global variance of temporal derivatives (GVTD)^31^. GVTD <u>sums the temporal derivatives of light levels over voxels as a parsimonious way of summarizing fluctuations in signal intensity,</u>conceptually similar to the DVARS measure sometimes used in fMRI^40,41^. <u>Previous studies have shown that GVTD is highly correlated with accelerometer-based measures of motion</u>^31^.

**Figure 3.**
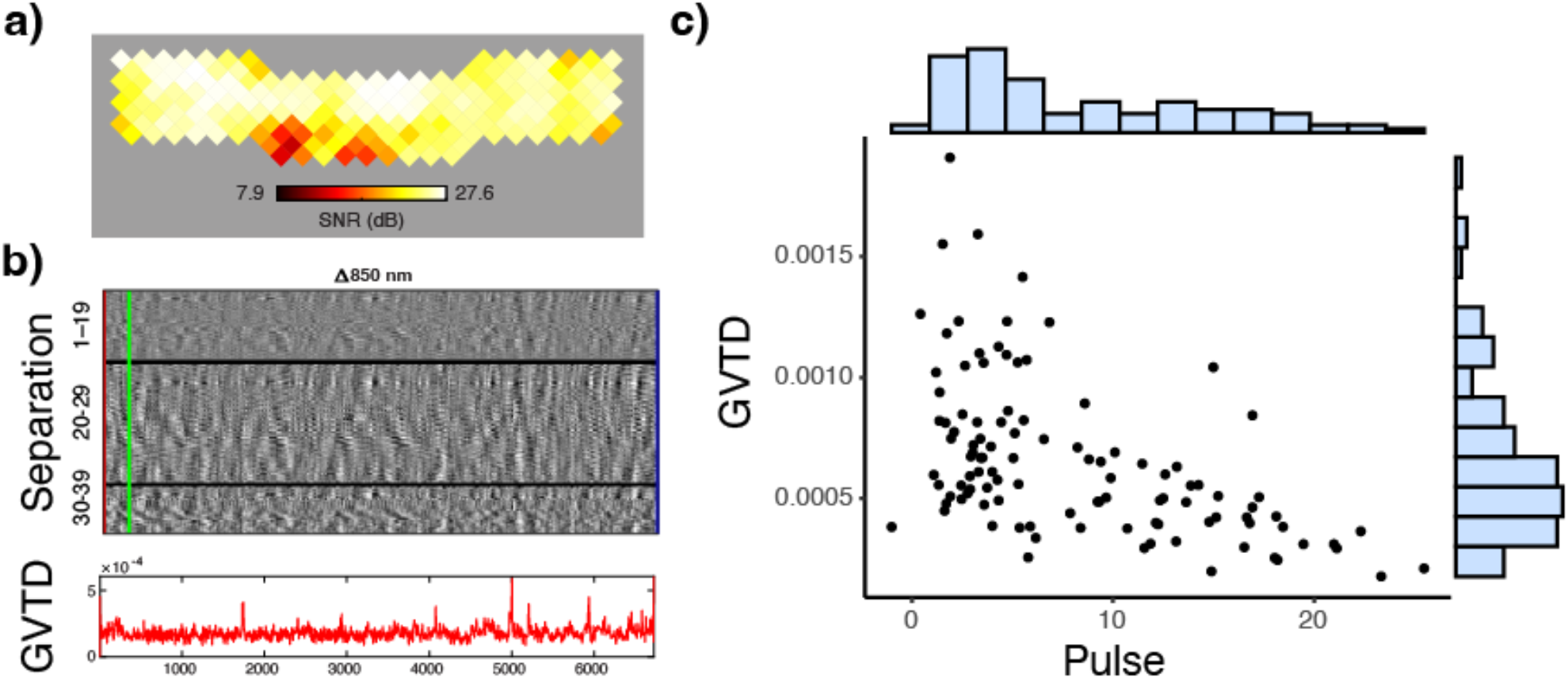
Quality control measures. **a)** Mean light levels over the cap for a sample participant (850 nm light). **b)** *Top:* example temporal derivative time traces from source detector distances between 1–19 mm, 20–29 mm, and 30–40 mm respectively, for 850 nm light in the same sample participant; *Bottom:* GVTD over the same time period. **c)** Scatter plot of GVTD (proxy for motion) and pulse SNR levels for all sessions.

We used the heartbeat (pulse) signal-to-noise ratio (SNR) as an indicator of detecting physiological signals in the data. <u>Data with low levels of pulse SNR are more likely to have low SNR levels for the hemodynamics frequencies as well. This is mainly due to a poor optode-scalp coupling.</u>The pulse SNR was calculated <u>using the NeuroDOT function</u>*<u>PlotCapPhysiologyPower</u>* as the proportion of the pulse power (based on the peak FFT magnitudes in the 0.5–2 Hz window) <u>of the optical density</u> signal divided by a noise floor (based on the FFT magnitudes in the 0.5–2 Hz window)^20^.

High quality data will contain low levels of GVTD and high levels of SNR. This is because GVTD is correlated with motion level (which is not desirable) and pulse SNR is correlated with hemodynamic signal SNR (which*is* desirable).

### Intersubject correlation analyses

In an intersubject correlation analysis, the similarity of time courses is computed for the same voxel in each pair of participants; averaging these values provides an average intersubject correlation value for every voxel of the brain^42^. Numerous studies have used intersubject correlation analysis to identify regions of the brain which show similar patterns of activity across a group of participants^43–45^. For movie viewing, these values are typically higher in sensory regions (e.g., visual cortex and auditory cortex) than regions associated with complex linguistic or executive processing^8,25^. One benefit of intersubject correlation analysis is that it is sensitive to shifts in the timing of activity across participants. Thus, demonstrating reasonable intersubject correlation values in sensory regions suggests the time courses across participants are correctly temporally aligned.

As part of our technical validation, we therefore performed an intersubject correlation analysis (**Figure 4a**). Because not all participants had multiple sessions of movie data, we restricted our analysis to the session with the highest pulse SNR from each participant. The signal average was subtracted from each frame and a Pearson correlation coefficient was then computed voxelwise using the corrected data for all pairs of subjects. We see the highest values in auditory and visual cortices, broadly consistent with prior studies in both fMRI^8^ and HD-DOT^28^.

**Figure 4.**
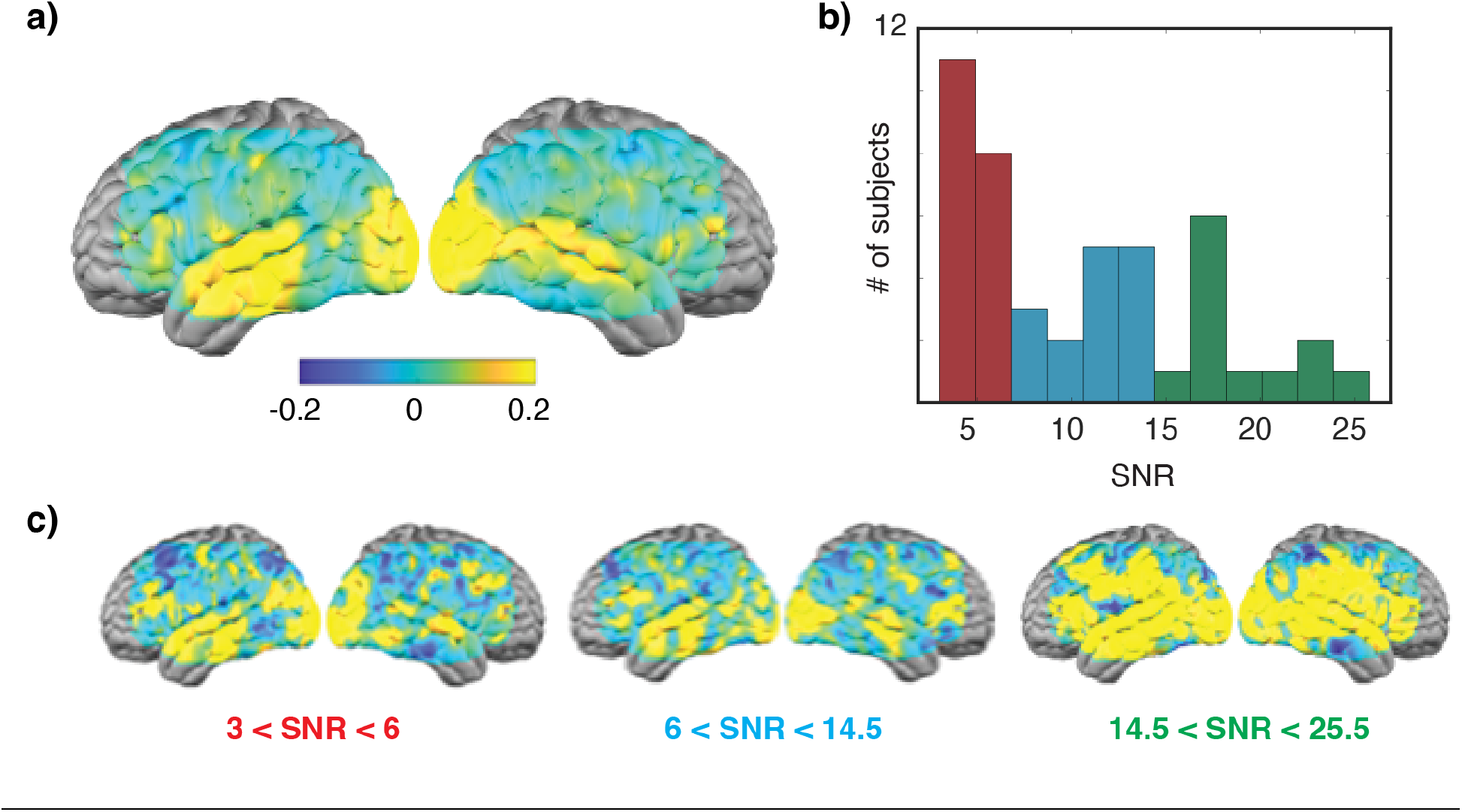
Intersubject correlation analysis. **a)** Correlation map shows the average pairwise correlation across all subjects. Maximum r values are observed in sensory areas. The SNR in these subjects, as quantified by pulse amplitude, ranged from approximately 3 to 25. **b)** A histogram of pulse values, and used to separate subjects into low (red), medium (blue), and high (green) SNR groups. **c)** Intersubject correlation maps obtained for subjects grouped by SNR. High correlations extend beyond sensory areas in the highest SNR group.

We then placed participants into one of three groups based on pulse amplitude measured in the optical signal (low group: 3.0 < pulse ≤ 6.0; med: 6.0 < pulse ≤ 14.5; high: 14.5 < pulse < 25.5) (**Figure 4b**). As shown in **Figure 4c**, we observed that r values were notably higher in subjects in the high pulse amplitude group compared to those in the medium and low groups, supporting our use of pulse amplitude as a quality control measure.

Finally, we wanted to make sure that the spatial pattern of our intersubject correlation maps was not a result of artifacts in the data. We adopted a censoring approach using GVTD to exclude frames with high variance (likely driven by motion). We used a GVTD threshold of 5E-04 to identify outliers. If the GVTD of a given frame exceeded this threshold in either subject during pairwise correlations, the frame was excluded when computing the correlation. Results from this analysis are shown in **Figure 5**. Although the absolute correlation values are reduced, we see a similar spatial pattern to the intersubject correlation maps, consistent with the data being correctly temporally aligned across participants.

**Figure 5.**
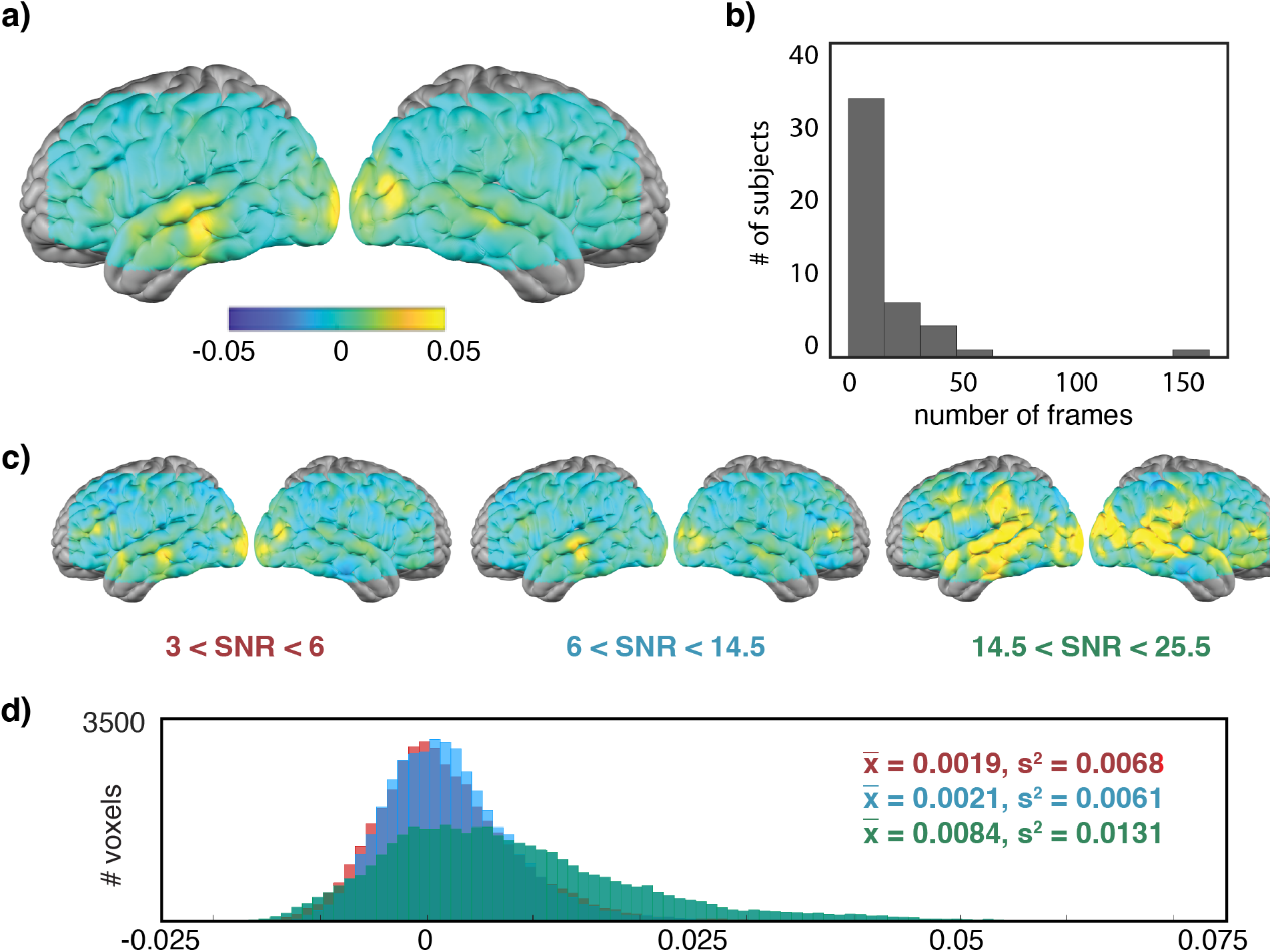
Results of intersubject correlation after adjusting for motion artifacts. Layout mirrors the organization of Figure 4. **a)** Correlation map shows the average pairwise Pearson correlation across all subjects. Maximum r values are observed in sensory areas. **b)** A histogram showing the number of participants who had different numbers of frames excluded for exceeding the GVTD threshold. **c)** Intersubject correlation maps on participants grouped by pulse SNR. (Pulse SNR was calculated prior to censoring, so these groups of participants are the same as those shown in Figure 4.) **<u>d)</u>** Histograms of voxel values in the three SNR groups. Mean and standard deviation shown at upper right. An average correlation value was calculated for all pairwise maps in each group and analyzed using Welsh’s ANOVA, which found a significant difference exists between the group means (F = 13.4, p=4.2E-06).

## Code Availability

Code used for technical validation is available from http://github.com/jpeelle/GBUDOT.

## Acknowledgments

This work was supported by R01 DC019507, R21 DC016086, R21 DC015884, R01 MH122751, K01 MH103594, R21 NS098020, and T32 EB014855 from the US National Institutes of Health. We thank Tessa G. George, Kelsey T. King, Karla M. Bergonzi, and Tracy M. Burns-Yocum for assistance with data collection.

## Author Contributions

J.E.P. conceived of the data sharing effort. A.F., A.S., A.B., M.M., and C.H.L. collected the data. E.S., A.B., and A.S. oversaw data conversion and organization. A.B., A.S., M.S.J. completed preprocessing and quality control analyses. M.S.J. conducted the intersubject correlation analyses. The manuscript was drafted by J.E.P., A.S., A.B., and M.S.J., and critically reviewed by all authors. J.E.P., J.P.C., A.T.E., and T.H. obtained funding.

